# Emerging Threat of Ranavirus: Prevalence, Genetic Diversity, and Climatic Drivers of *Ranavirus* (Family Iridoviridae) in ectothermic vertebrates of Asia

**DOI:** 10.1101/2023.08.19.553946

**Authors:** Jayampathi Herath, Sun Dan, Gajaba Ellepola, Kuttichantran Subramaniam, Madhava Meegaskumbura

**Author notes:** These authors contributed equally (Jayampathi Herath, Sun Dan). Corresponding author. E–mail.

## Abstract

Ranavirus disease, caused by viruses within the genus *Ranavirus* (family *Iridoviridae*), is considered a globally emerging infectious disease linked to mass mortality events in both wild and cultured ectothermic vertebrates. Surveillance work is however limited in Asia hence prevalence and the dynamics of the disease remains poorly understood. To understand disease burden and the potential biotic and abiotic drivers in southern China region, we conducted a systematic surveillance of the ranavirus across Guangxi Zhuang Autonomous region (GAR). For this, we used a multifaceted approach involving screening of amphibians and other potential reservoirs, diagnostic tests, phylogenetic analyses, prevalence estimation, co-infection assessments, and climatic niche analyses. Over one thousand individuals were sampled across 25 sampling sites. We found ninety-two individuals from 18 species of ectothermic vertebrates to be infected with ranavirus. Two lineages were responsible – Rana nigromaculata ranavirus and Tiger frog virus were identified using phylogenetic analysis based on the major capsid protein (MCP) gene fragment. We also found evidence of a co-infection with ranavirus and *Bd* that can be highly detrimental to host populations; possibly the first such documentation in Asia. Our niche modelling analysis suggests that precipitation and seasonality play an important role in ranavirus prevalence in Guangxi region – southwestern, southeastern, central and northeastern regions of GAR can be considered to be optimum habitats for ranaviruses. Infection rates in wild frog species have reached 100% in some areas, even in nature reserves. Our research also indicates that culture facilities and pet farms are frequently infected, serving as likely vectors for the regional and global spread of ranaviruses. The knowledge generated suggests the need for systematic surveillance, stringent biosecurity measures, and control of international animal trade to prevent further transmission and protection of biodiversity and aquaculture industries across Asia.

## INTRODUCTION

Ranaviruses, a group of double-stranded DNA viruses within the genus *Ramavirus* (family *Iridoviridae*; subfamily *Alphairidovirinae*), have emerged as a major threat to amphibian populations worldwide, as well as other ectothermic vertebrates (Gray and Chinchar, 2015). Thirty-five viruses belonging to 15 viral families (11 with RNA genomes and 4 with DNA genomes) which are infecting amphibians and reptiles have been identified (Harding *et al*., 2022). Ranaviruses have been linked to mass mortality events and are contributing to the ongoing decline of amphibians along with fish and reptile populations (Chinchar and Waltzek, 2014), heightening concerns about biodiversity loss and ecosystem functionality. Ranaviruses are thought to be a new arrival in Asia, with relatively unknown effects on biodiversity (Herath *et al.,* 2021). As a region with exceptional amphibian diversity and habitats under anthropogenic pressure, Asia presents an important region for investigating the occurrence, distribution, and potential drivers of ranavirus infections.

Ranaviruses are known to infect a broad host range, which adds to their potential for causing widespread ecological damage. The virulence of these pathogens can be exacerbated due to various factors such as environmental conditions, seasonality, and host density and immune responses (Brunner *et al.,* 2015). Ranaviruses are typically transmitted through direct contact between individuals, ingestion of infected tissues, or contact with contaminated water or fomites (Bruner *et al*., 2015; Miller *et al*., 2011). This mode of transmission enables the rapid spread of the virus within populations, leading to high morbidity and mortality. Currently, there are seven species within the genus *Ranavirus*, with three potential new species remaining unclassified (Chinchar *et al*., 2017). Frog virus 3 (FV3) and related ranaviruses primarily infect amphibians as well as fish and reptiles, while Ambystoma tigrinum virus (ATV) predominantly affects caudate amphibian specialist and few studies have shown that some anurans are susceptible to the disease as well. The most recent common ancestor of common midwife toad virus (CMTV), FV3, and other closely related ranaviruses appear to have infected amphibians. However, CMTV may circulate independently within both amphibian and fish populations (Price *et al*., 2017). As such, understanding the host-specificity, ecology and epidemiology of ranaviruses in Asia is crucial for predicting and managing their impacts on regional biodiversity, aquaculture, and ecosystem functioning.

It is also essential to understand the impact of ranaviruses on amphibian populations in Asia for several reasons. First, many Asian amphibian species are already threatened by habitat loss, pollution, and overexploitation (Rowley *et al*., 2010). These factors may make them even more vulnerable to ranavirus infections (Brunner *et al*., 2015). Identifying the presence and prevalence of ranaviruses in the region can inform targeted conservation efforts and help prioritize resources to protect the most vulnerable species and habitats.

Asia also serves as a nexus for global trade and wildlife trafficking, resulting in importing of a large number of species for both consumption and the pet industry (Hughes, 2021; Kolby *et al*., 2014). This increases the risk of ranavirus spreading through the inadvertent movement of infected specimens (Herath *et al*., 2021). The ability of ranavirus to be transmitted through water among all ectothermic vertebrate classes makes it one of the highly transmissible diseases (Brenes *et al*., 2014). Investigating the genetic diversity and relationships among ranaviruses in Asia can thus provide insights into their origins, transmission pathways, and evolutionary patterns, enabling the development of targeted interventions to prevent further spread.

The diverse climate and topography of southern China provide an opportunity to examine the potential drivers of infection prevalence, such as bioclimatic variables, elevation, season, and habitat-related factors. Understanding these drivers is also critical for developing targeted strategies to reduce the spread of ranaviruses and mitigate their impact on amphibian populations.

Finally, studying co-infections with other pathogens affecting amphibians, such as *Batrachochytrium dendrobatidis* (*Bd*), can offer a deeper understanding of disease dynamics in the region. This will help inform integrated disease management strategies. By examining the occurrence, distribution, and drivers of ranavirus infections in Asia, we aim to contribute to the global effort to protect amphibian populations and conservation of biodiversity.

In the current study, we investigate the occurrence of ranaviruses in amphibian and other ectothermic vertebrate populations in southern China and assess the potential drivers of infection prevalence of the diseases, thus generating vital knowledge for conservation. For this, we will analyze the ranavirus infection across a vast swath of land in southern China with an emphasis on the following specific points: (1) Conduct diagnostic tests, to identify the presence of ranavirus in amphibians and potential reservoir hosts (2) Carryout genetic analysis based on the major capsid protein (MCP) gene regions to determine the ranaviruses lineages and their phylogenetic relationships. (3) Estimate the infection prevalence (4) Assess co-infection with *Bd* in amphibian samples. (5) Analyze the climatic niche to understand the potential distribution of ranaviruses in the southern China region and identify the environmental factors that contribute to their occurrence. Investigating the occurrence of ranavirus and potential drivers of infection in amphibian populations of southern China is an important step in preventing the regional and global spread of these diseases.

## MATERIALS AND METHODS

### 1. Identifying the presence of ranavirus in southern China

We carried out field surveys and diagnostic tests, to uncover and monitor whether it is a disease hotspot.

#### Field sites

Guangxi Autonomous Region (GAR), situated in southern China is bordered by Yunnan to the west, Guizhou to the north, Hunan to the northeast, Guangdong to the east and southeast, gulf of Tonkin in the south and Vietnam in the southwest (21°42.45’-25°37.01’ N, 107°32.59’-110°12.44’ E). This massive plain covers an area of 237,600 km^2^, with some mountainous terrain. Several river systems including Qin and the Nanliu Rivers flow into the Gulf of Tonkin. Several tributaries flow into the larger Xiang River in neighboring Hunan province, and the Xi River system flows southeast. This subtropical region is moist and warm with an annual precipitation ranging from 723.9∼2983.8 mm, and the annual mean temperature is between 17.6∼23.8 °C (Hao *et al*., 2023). The region receives substantial precipitation during the monsoons arriving from south-southwest in late April to the beginning of October. Unique karst landforms are found with the central parts forming a basin surrounded by areas of higher elevation (Hao *et al*., 2023).

#### Main sampling sites

We sampled across a large swath of land in GAR representative of the environmental heterogeneity of south China. Sampling sites covered a wide range of altitudes and vegetation types including sub-tropical evergreen broad-leaved forests in the north and sub-tropical evergreen seasonal rainforests in the south. The sampling design comprised of paired sampling sites: one within nature reserves representing undisturbed habitats and another outside the nature reserve representing disturbed habitat within average distance of 10 km. A total of 12 main sampling sites were selected, namely: Shiwan Dashan, Nakuan, Pinglong, Dongzhong, Dayaoshan, Cenwanglaoshan, Shengtangshan, Hongtan, Anjiangping, Wuzhishan, Daling and Cujiang (Fig: 1). Sampling was carried out from September 2018 to September 2021 capturing a representation of the seasonal changes as well (Fig: 1).

#### Sporadic sampling

Sporadic sampling was carried out in several locations in Guangxi region, including both natural and disturbed habitats. These included ponds, paddy fields, streams and seasonal ponds, river and home gardens (Fig: 1).

#### Culturing facilities, Markets and Pet markets

In addition to major sampling sites, we carried out continuous and opportunistic sampling in markets and pet markets, where numerous indigenous and exotic ectothermic vertebrate species were present. Many of these animals were housed under substandard conditions and imported from countries with known ranavirus infections. These facilities pose a high risk for disease spread as some of the species are known host species of ranavirus infection. It is essential to screen for the disease in these locations to understand the potential for disease transmission from captive to wild animals. Swabs were taken from fish (both freshwater and marine), turtles and frogs that were sold at several markets across various locations. Frog farms included Luobo, Yulin and Bobai frog farms, while markets included Guigang, Luobo, Gunagxi University, Jin Xiu County and Fangchenggang seafood markets together with two supermarkets in Nanning. Further, sampling was also done in Nanning pet market, Guangxi pet market, Guigang pet market and on several specimens from several private owners (Fig: 1).

#### Sampling

We sampled potential hosts from various parts of the forest including leaf litter, floor, trees and various water bodies, both periodic and perennial, such as rivers, streams, lakes and ponds. The primary focus was on sampling amphibians, including their egg clusters and larval stages, as well as fish and reptiles. A small number of aquatic mollusks, shrimps, and crabs were also sampled to investigate their potential as carriers. Hand nets, umbrella nets and traps were used for sampling. Both diurnal and night sampling were carried out to represent all the species living in that natural habitat. Further, randomly selected specimens representing all the ectothermic vertebrates were sampled in frog farms, markets and pet markets with prior approval from the owners. Individuals were temporarily kept in new clean and unused 10 x 5 cm or 15 x 20 cm plastic zippered bags with a few holes punctured for ventilation. All the species were photographed and identified to the species level using standard guides. Data was recorded along with GPS coordinates, and photographs, including the species and life stage.

#### Swabbing

We used non-lethal swabbing for sample collection following Gray *et al*., (2012). Sterile, dry swabs with fine tips (Medical Wire & Equipment Co. MW 113) and plastic shafts (to avoid PCR inhibitors) were used. Swabbing was performed gently but firmly, swiping the swab along the surface to be tested. Surfaces typically swabbed for ranaviruses include the oral cavity, cloaca, or skin lesions (Pessier and Mendelson, 2017). Swabbing the vent provides evidence of intestinal shedding and swabbing the cloaca offers a high chance of capturing internal viral shedding. Immediately after swabbing, the shaft was broken and swabs were put into 1.5 ml Eppendorf tubes without touching, before releasing the amphibians back to their point of capture. The swabs in Eppendorf tubes were stored at −80°C in the lab until DNA extraction. In the event of dead were encountered, symptoms were recorded, and swabs were taken from internal organs such as the liver, when possible, primarily in frog farms.

#### Avoiding cross contamination

To avoid cross-contamination, powder free nitrile disposable gloves were worn, and these were changed between each individual sampled. Individual animals were not co-housed; they were captured and stored in plastic bags individually until swabbed and released. For tail or toe clips, sterile instruments were used to avoid sample contamination. A 4 % bleach solution was used for inactivating ranavirus and other pathogens, such as the amphibian chytrid fungus (Bryan *et al*.,2009; Gold *et al*., 2013) when cleaning the sampling utensils. Boots, waders, nets, traps, clothing, or other equipment that was exposed to water or mud were thoroughly washed once surveys were completed at each site to remove any lingering mud containing pathogens. They were then decontaminated using a mixture of 10% bleach (Pessier and Mendelson, 2017).

#### Ethical clearance

Ethical clearance was obtained from the Institutional Animal Care and Use Committee of Guangxi University (GXU2018-048, with the extension of GXU2020-501). All the procedures were carried out according to the standard ethical practices and protocols while no animal was sacrificed. Prior permission was obtained from Nature reserves and protected areas; relevant regulations and protocols were followed.

#### DNA extraction

QIAamp UCP Pathogen Mini Kit was used for DNA extractions which were used for Real Time PCR (RT-q PCR). Protocol of pretreatment of Microbial DNA from Eye, Nasal, Pharyngeal, or other Swabs (without Pre-lysis) was used. Qiagen DNeasy Blood and Tissue Kit was used for DNA extractions following guidance of the producer which were used for conventional PCR.

#### Quantitative Real Time PCR (RT-qPCR)

Initially, real-time quantitative PCR (RT-qPCR) was employed for disease surveillance in the collected swabs from GAR during the preliminary stage of the sampling process. Preliminary surveillance was essential since the preliminary data was not available. Prior to this study, only one outbreak in cultured hybrid grouper had been recorded in Guangxi (Xiao *et al*., 2019). The RT-qPCR was used because it is known to be more sensitive than conventional PCR with the ability to detect lower viral loads. qPCR primers (RanaF1 5’-CCA GCC TGG TGT ACG AAA ACA −3 and RanaR1 5’-ACT GGG ATG GAG GTG GCA TA −3’) TaqMan probe (RanaP1 6FAM-TGG GAG TCG AGT ACT AC-MGB) targeting a conserved region of the major capsid protein (MCP) gene (Stilwell *et al*., 2018) were used for 60 samples. This assay has been designed to detect a panel of 33 different ranaviral isolates originating from fish, amphibian, and reptile hosts, representing the global diversity of ranaviruses. Roche LightCycler® 480 was used for the RT-qPCR work. Amplification conditions were set as follows: an activating cycle at 95℃ (10 min), and then 45 cycles at 95 °C (10s), 60 °C (10s) and 70 °C (1 s) followed by a cycle 40 °C (30 s). All plates were run with a negative control (nuclease-free water) and a known positive. Positive samples were run again in the same machine and only samples with consecutive positive results were declared positive. Subsequently, conventional PCR followed by Sanger sequencing were performed as the next stage of analysis.

#### PCR and gene sequencing

PCR was performed using primers (forward primer: 5′-GACTTGGCCACTTATGAC-3′ and reverse primer: 5′-GTCTCTGGAGAAGAAGAA-3′) targeting highly conserved regions of the MCP gene (Mao *et al*., 1997). The PCR conditions consisted of 4 min at 94°C, then 35 cycles of 30 s at 94°C, 30 s at 55°C, and 1 min at 72°C, followed by 10 min at 72°C. Negative and positive controls were included in each PCR amplification. Amplicons of the expected size (500 bp) were purified and sent to a commercial Sanger sequencing service (Sangon Biotech, Shanghai). A total of 78 samples were sequenced.

### 2. Phylogenetic analysis based on the major capsid protein (MCP) gene

We conducted a phylogenetic analysis to establish relationships among identified ranaviruses, offering insights into their origins and transmission pathways. Phylogenetic inference was based on the MCP gene fragment, utilizing sequenced samples from this study combined with sequences of ranaviruses obtained from the GenBank. The MCP gene sequences were aligned using MUSCLE, as implemented in MEGA v.6.0 (Tamura *et al.,* 2013). Regions with low confidence in positional homology were removed from the analysis and edges were trimmed. Prior to constructing the phylogenetic tree, the best-fitting nucleotide substitution model was determined using jModelTest v.2.1.4 (Darriba et al. 2012; Guindon and Gascuel, 2003). A maximum likelihood (ML) tree was built using MrBayes and MEGA, while a Bayesian tree was constructed using BEAST and an IQ tree in Phylosuite was constructed, all of which yielded similar topologies; only the ML tree is presented here (other trees are provided in supplementary materials)

### 3. Estimating the infection prevalence across sampling sites Infection prevalence

We estimated infection prevalence to identify areas and species at risk that will enable focused conservation efforts. Infection prevalence is a measure that estimates the proportion of a population infected at a specific point in time (Gray *et al.,* 2015). This measure can be regarded as a “snapshot” of the infection burden at a given time. Infection prevalence was determined for positive cases recorded in main sampling sites, sporadic sampling sites, and frog farms. However, it was not calculated for markets and pet markets, as these are temporary holding facilities, and the animals originate from various locations.

### 4. Investigating the occurrence of co-infection with ranavirus and *Bd*

We assessed co-infections with *Bd* to provide a comprehensive understanding of disease dynamics and interactions between pathogens that will inform integrated disease management strategies.

Swabs obtained from the main sampling sites; Shengtangshan, Hongtan, Nakuan, Anjiangping, Pinglong, Hongtan, Dongzhong, Wuzhishan, Daling and Cujiang were tested for *Bd* as well (as a part of another ongoing study) (n = 501). A nested PCR assay for the detection of *Bd* was carried out (Annis *et al*., 2006; Goka *et al*., 2009).

### 5. Climatic niche and distribution analyses

Finally, we analyzed the climatic niches of ranaviruses to gain insights into their potential future distribution under changing environmental conditions and inform proactive measures to prevent the spread of ranaviruses and protect vulnerable amphibian populations.

#### Drivers of infection prevalence analysis

The prevalence of ranavirus was assessed using generalized linear models (GLMs). Information on 19 bioclimatic variables for each occurrence point was obtained from WORLDCLIM 2.172 using the “extract” function in RASTER 4.2.2.

An information-theoretic modelling approach was performed (Burnham, 2002) to assess the effects of multiple bioclimatic, elevation, season (month), habitat factors, and life history. Nineteen bioclimatic variables were downloaded at a resolution of 30 arc sec (Stephen and Hijmans, 2017). We calculated the correlation between bioclimatic factors, and only selected 4 variables (bio7, bio8, bio15 and bio18) with a correlation coefficient < 0.70 for further analysis. To classify adult habitats, we used the activity breadth of adults observed during the non-breeding season (Laurent, 1964; Bardua *et al*., 2021).

A Generalized Linear Model (GLM) was constructed, as a candidate model based on all possible combinations, to analyze the influence of eight predictor variables (life history, bio7, bio8, bio15, bio18, elevation, habitat, month) on ranaviruses prevalence (infected individuals/all individuals) in main sampling sites, sporadic sampling sites and frog farms. Meat markets and pet markets were not considered as temporary holding facilities where the animals are kept for short time periods. These variables were set as explanatory variables, as well as together with a null model. We used populations infected by ranaviruses within sites as the response variable. were set as explanatory variables, as well as together with a null model. We also included a candidate model with species as the single explanatory variable was also included to assess whether species per se affect the infected individuals/all individuals.

Each candidate model was quantified and evaluated based on Akaike Information Criterion (AIC), Akaike second order corrected (AICc) and Akaike weights (AICw) (Steidl, 2008). The final support model was validated according to the evaluation of homogeneity in the residuals of the models against fitted values (Zuur *et al*., 2010).

#### Species distribution modeling using MaxEnt

We used presence and absent data obtained during the current study to build a species distribution model for ranaviruses in the Guangxi region. We included 19 presence localities and 518 absent localities of Ranaviruses in the Maxent species distribution model that we constructed. A biased grid was used for data thinning and reducing sampling bias. Information on 19 bioclimatic variables for each occurrence point was obtained for present day conditions (∼1970-2000) from WORLDCLIM 2.172 using the “extract” function in RASTER 3.0-773 at a spatial resolution of 30 arcsecs (∼1km2). Predictor collinearity was eliminated by calculating Pearsons’s correlation coefficients for all pairs of bioclimatic variables, excluding the variables from a correlated pair (|r| > 0.85. After excluding the correlated variables, bio2, bio3, bio5, bio11, bio16, bio17 and bio18 were used to build the model. The MaxEnt model was optimized using the ENMeval package (Muscarella *et al.,* 2014) by looking for the best AUC value after assigning a range of regularization coefficient values (0.05, 0.95, 0.05) for linear and quadratic features by looping the code. A random number generator was used for selecting 70% of the presence data for model building and the other 30% of presence data was used for evaluating the model (model test). Model performance was measured using the Area Under the Curve (AUC) and the results were overlaid on raster maps.

#### Land use factors on ranavirus presence

The spatial data map of the Guangxi region containing land use patterns was overlaid on the constructed niche model. Inferences were made based on the 13 available spatial data categories.

## RESULTS

### Presence of the ranaviruses in Guangxi Zhuang Autonomous Region

#### Epidemiology of ranavirus

In total, 1076 individuals from various sampling sites were examined, with 92 infected individuals identified across 18 species of ectothermic vertebrates. These included 84 PCR-positive and 8 qPCR-positive cases, encompassing 14 amphibian species (13 anurans and 1 caudate), 3 fish species, and 1 reptile species (testudine) (Fig 2 a, Table 1). The infected species were from natural environments and culture facilities throughout the Guangxi region (Table 1). Anurans from the Dicroglossidae family exhibited the highest mean infection prevalence, followed by Rhacophoridae, Ranidae, Microhylidae, and Hylidae (Fig. 2 c). The majority of infected ectothermic vertebrates were aquatic (12 species), with 4 species being semiaquatic and 1 species each from terrestrial and arboreal habitats (Fig. 2 b). A freshwater crab species tested positive for infection at Laohuling, likely due to environmental contamination from three infected frog species inhabiting the same pool.

**Figure 1:**
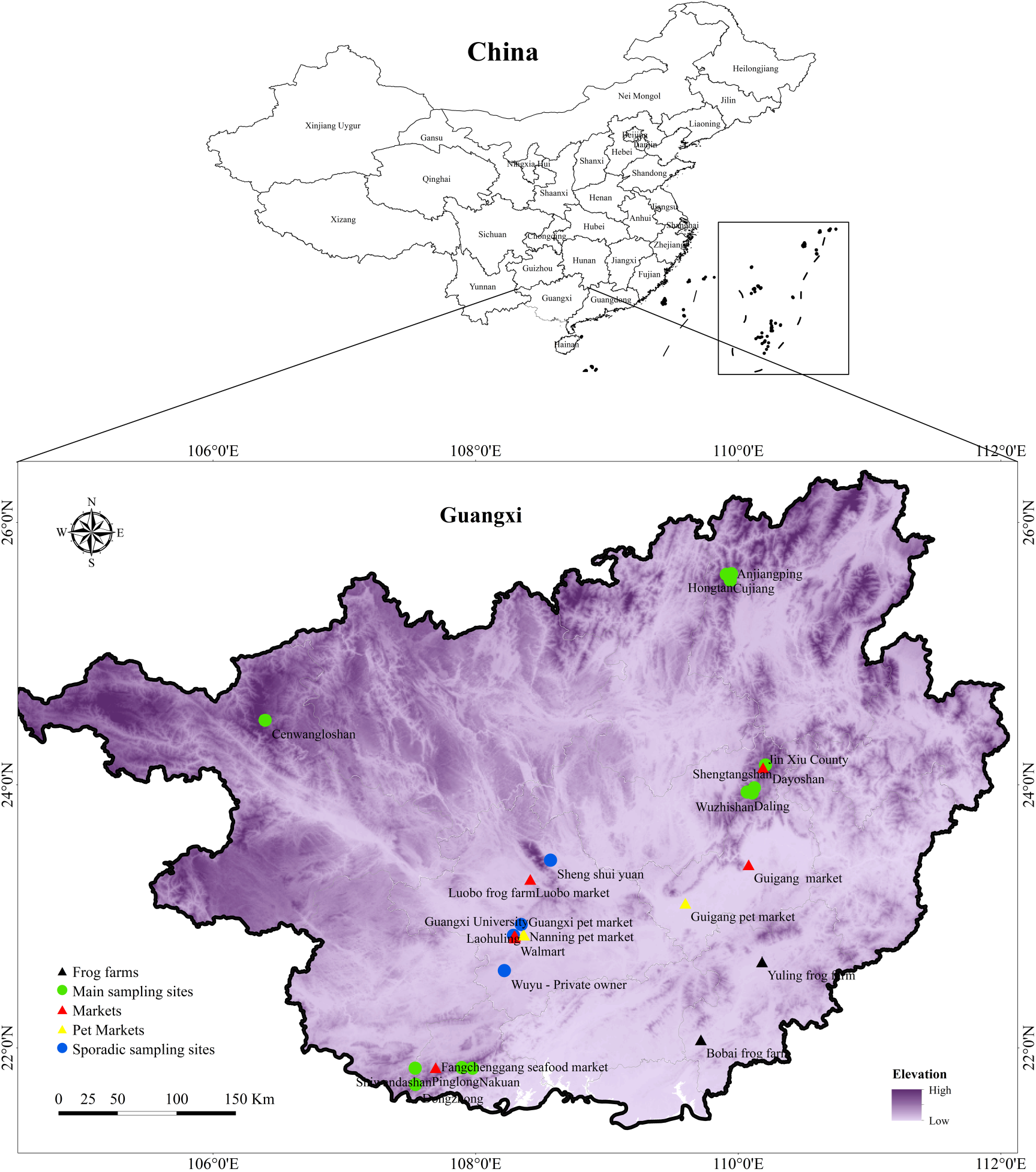
Map of sampling sites. The map depicts the distribution of main sampling sites, sporadic sampling sites, frog farms, markets and pet markets across GAR.

**Figure 2:**
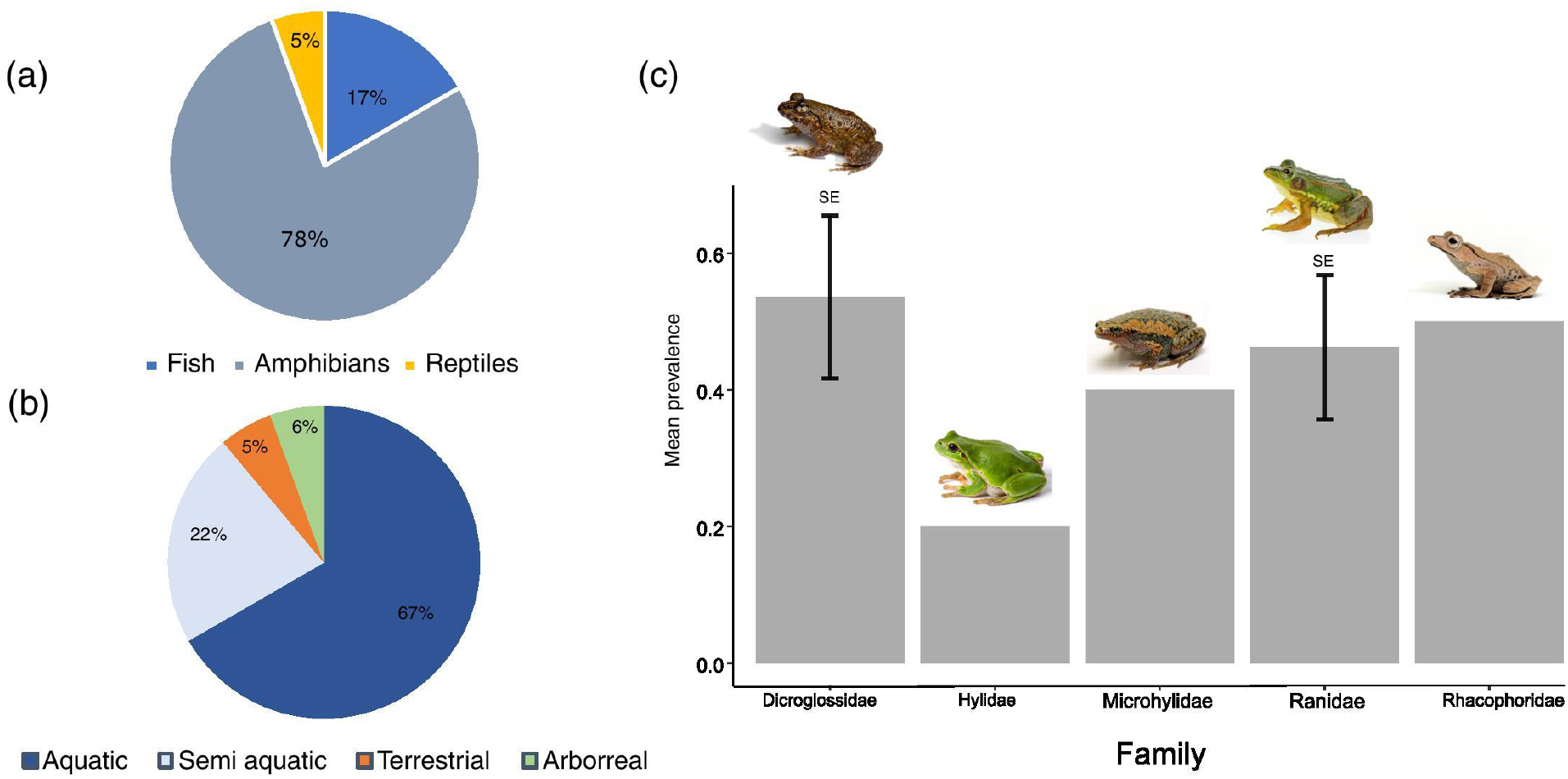
Epidemiological characteristics of the disease. (a) Number of species infected according to class. (b) Number of species infected according to their habitat. (c) Mean infection prevalence percentages among different families of anurans. Standard error is only presented for Families which have more than 1 species. Ranidae (n = 7 species), Dicroglossidae (n = 3 species), Rhacophoridae (n = 1 species), Microhylidae (n = 1 species), Hylidae (n = 1 species).

**Table 1:**
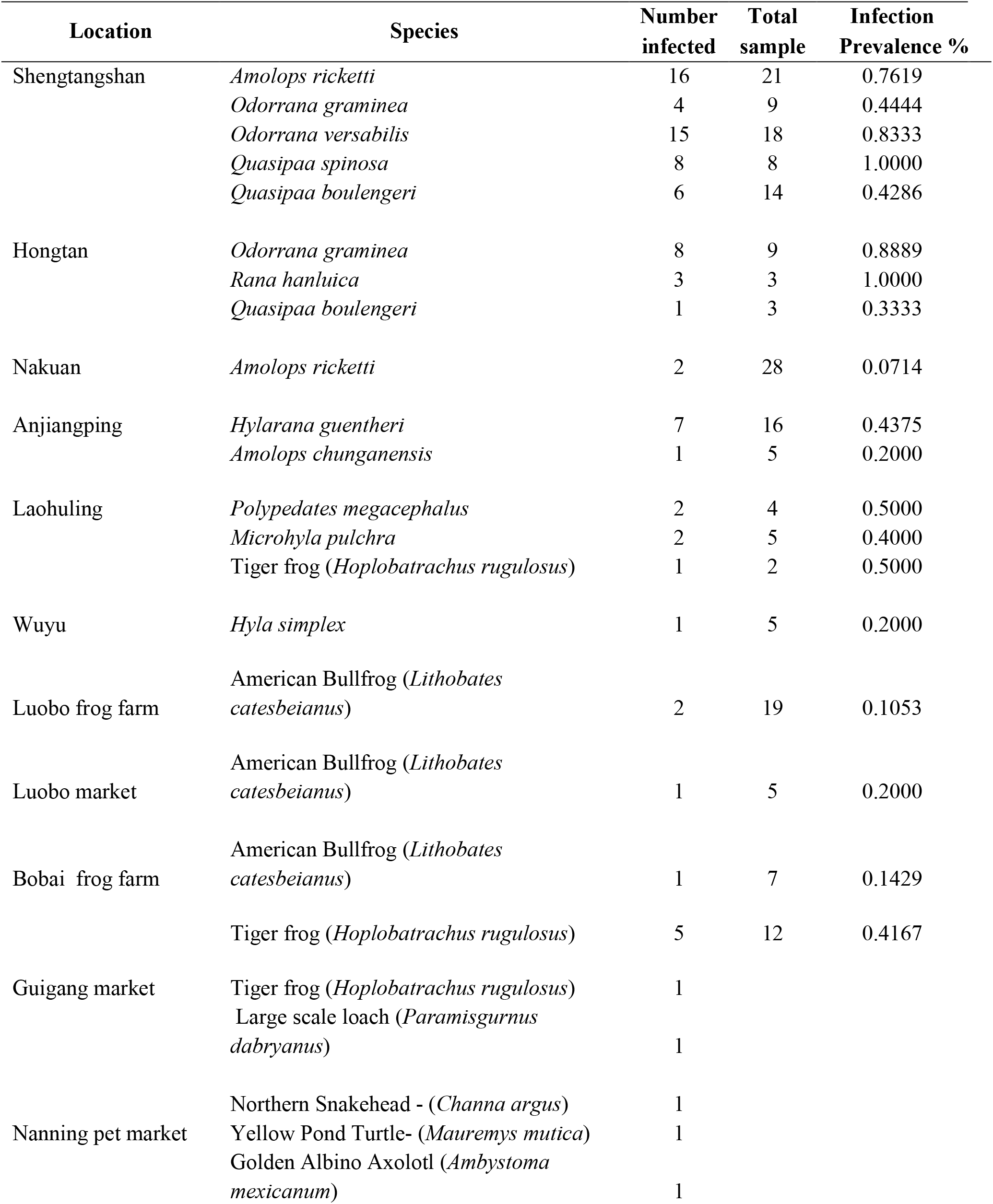
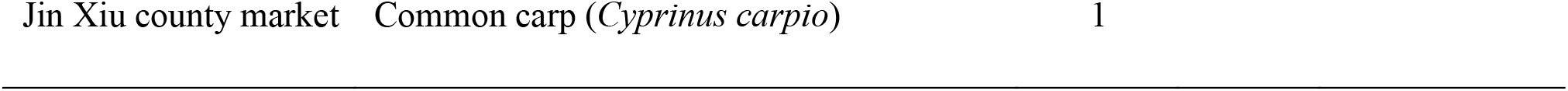
Infection prevalence in GAR Represents the sampling sites, host species, number of infected individuals of a species per a site, number of total sampled individuals of a species per a site, infection prevalence and infection prevalence percentage.

Most infections were detected during summer and were found in both adult and larval stages of anurans. In the majority of cases, there were no notable disease symptoms or mortality.

#### Clinical and behavioral signs

Cutaneous ulceration and hemorrhages are two common clinical signs in frogs associated with ranavirus infection (Cunningham et al., 2007). Cutaneous ulceration was visible on the infected yellow pond turtle from the Nanning pet market (Fig 3a) and the moribund large-scale loach from the Guigang market (Fig 3c). Erythema and ulcers (Fig 3b) with erratic swimming behavior and loss of equilibrium were observed on the infected tiger frog from the Bobai frog farm (Fig 4)

**Fig 3:**
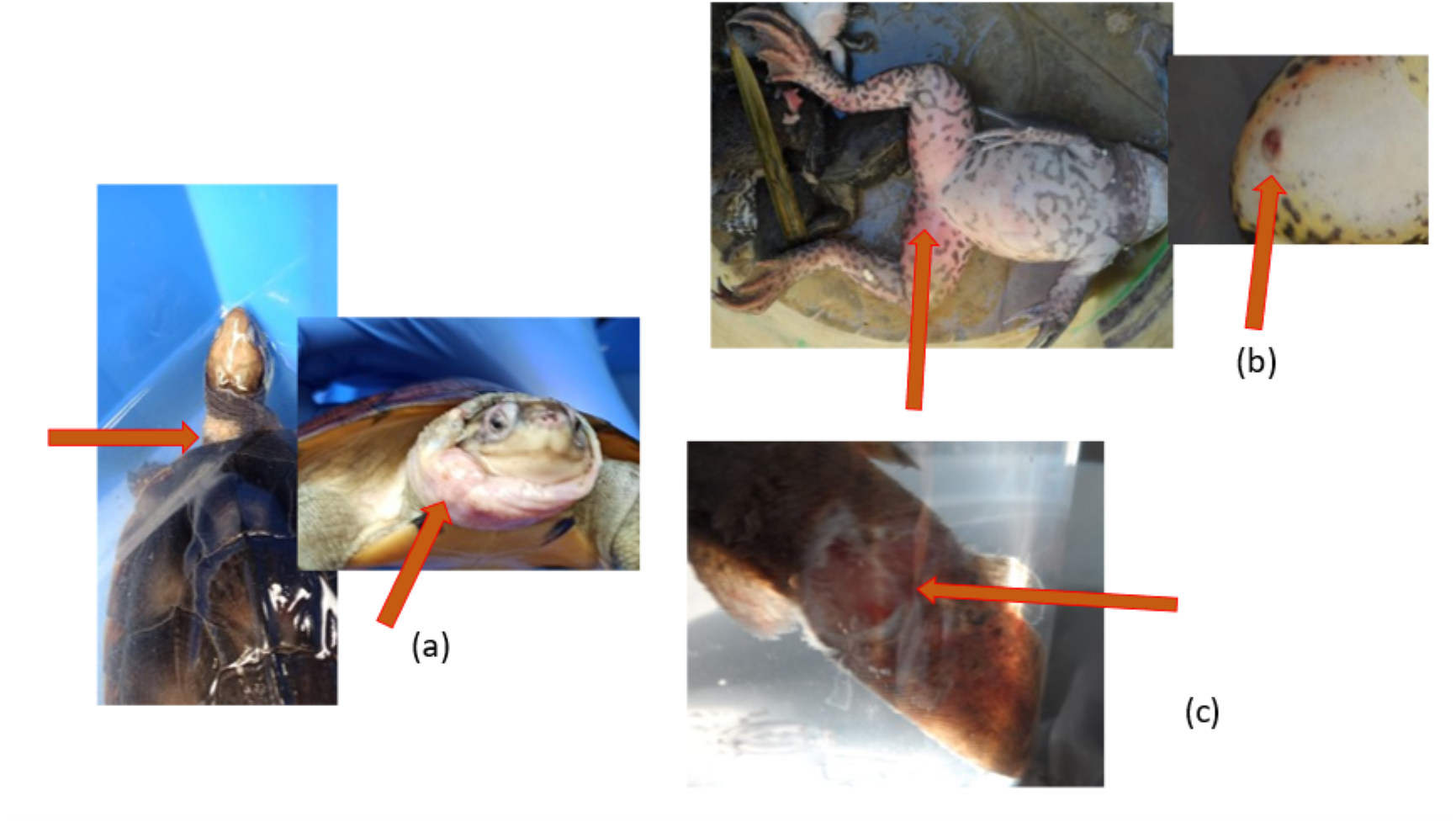
Gross lesions associated with ranavirus infection. (a) Yellow pond turtle (*Mauremys mutica*) displaying cutaneous ulceration on the neck and head. (b) Tiger frog (*Hoplobatrachus rugulosus*) displaying erythema and cutaneous ulceration on the ventral aspect of the hindlimbs. (c) Large scale loach (*Paramisguruns dadryanus*) displaying cutaneous ulceration of the caudal peduncle.

**Figure 4:**
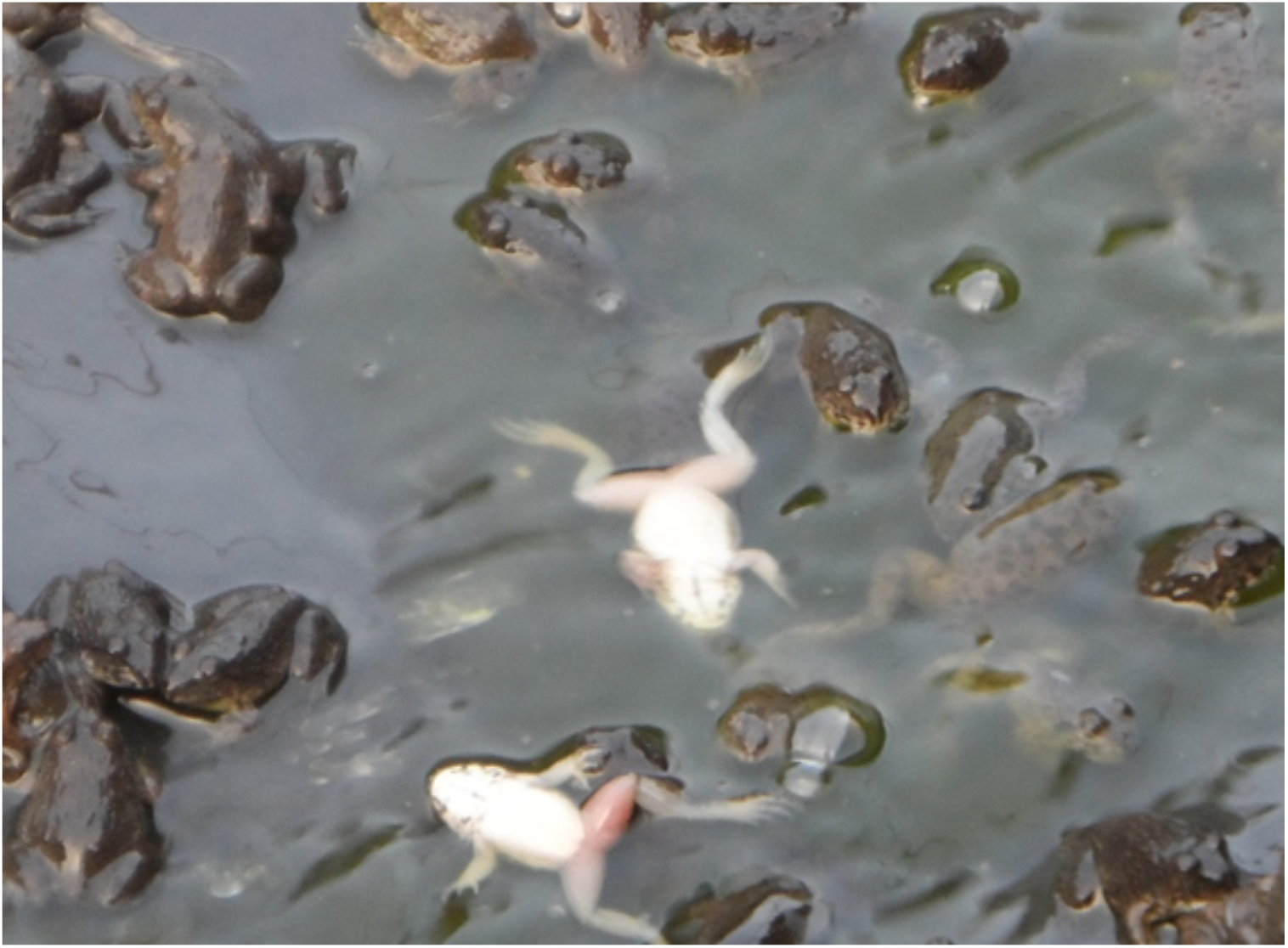
Erratic swimming behavior observed in tiger frogs.

### 2. Phylogenetic analysis based on the major capsid protein (MCP) gene to determine the genetic relationships

All sequenced samples were identical to Rana nigromaculata ranavirus (as the pairwise distance between the sequences was zero) except for one sample resembling tiger frog virus. Rana nigromaculata ranavirus was found in all the positive species in Shengtangshan, Hongtan, Nakuan, Anjiangping, Laohuling and Wuyu as well as two *Polypedates megacephalus* from Laohuling, two *Microhyla pulchra* from Laohuling, one *Hyla simplex* from Wuyu and one tiger frog from Bobai frog farm. The only detection of tiger frog virus was recorded from a tiger frog originated in Bobai frog farm (Fig 5).

**Figure 5:**
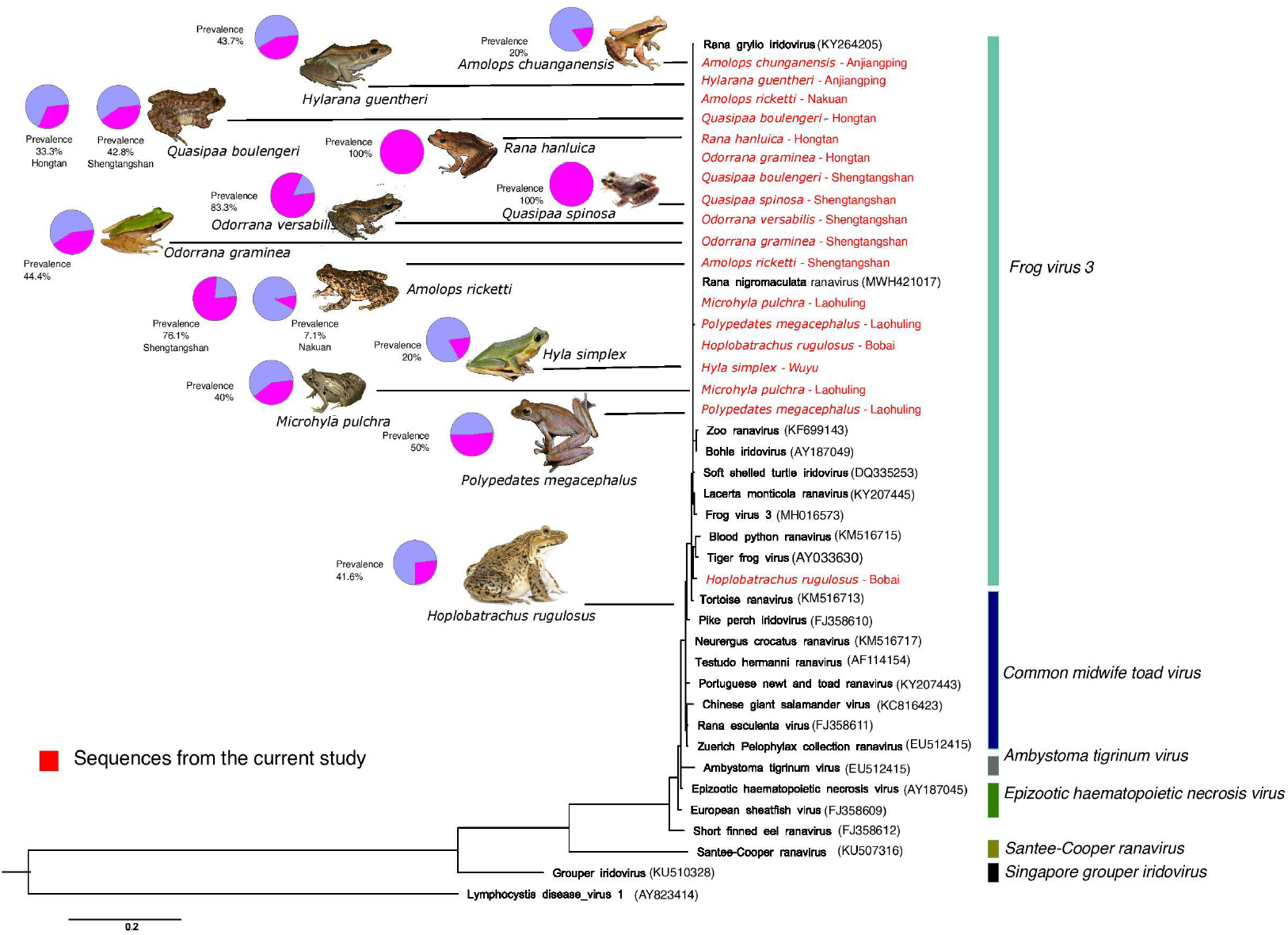
Phylogenetic tree based on the MCP gene sequenced. Highlighted in red are the sequences produced from this study. *Only 18 samples out of 77 are shown here. Seventy-six sequences are identical to Rana nigromaculata ranavirus and each other.

### 3. Estimate the infection prevalence in main sampling sites, sporadic sampling sites, culturing facilities, markets and pet markets

#### Disease prevalence

Highest infection prevalence percentages (100%) were observed from *Q. spinosa* from Shengtangshan and *R. hanluica* from Hongtan (Table 1). Both of these cases were recorded from natural habitats and exhibited higher prevalent rates compared to sporadic sampling sites and Culturing facilities.

#### Location prevalence

Infections were detected in 12 out of 29 sampling sites, with a location prevalence of 41.37%. Among the main sites, infections were recorded in 4 out of 12, while 2 out of 4 sporadic sampling sites had infections. Frog farms had a high infection rate (2/3), with infections found in 3 out of 7 markets and 1 out of 3 pet markets (Fig 6).

**Figure 6:**
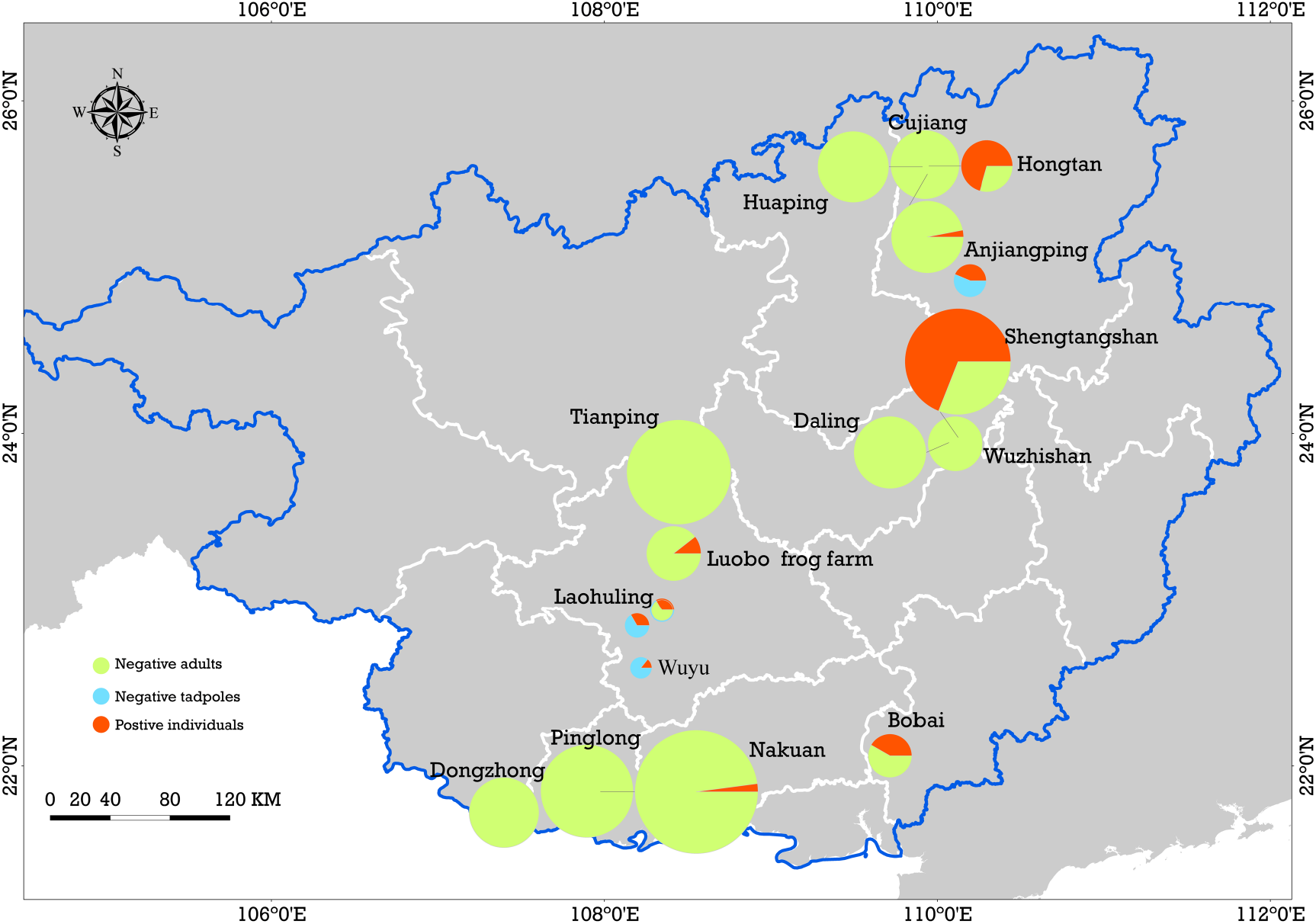
Map of sampling sites and infection prevalence recorded. Represents the Infection prevalence of main sampling sites, sporadic sampling sites and frog farms. The size variations of the circles correspond to the numbers of skin swab samples.

### 4. Occurrence of co-infection with ranavirus and *Bd*

Two frog species, *O. graminea* (LC IUCN status) and *Q. boulengeri*, (VU IUCN status) were found to have co-infections of both ranaviruses and *Bd* out of 501 samples. Co-infections are present in GAR though the rate of co-infection is very low. There were no records of *Bsal* found in GAR. (Sun *et al*., 2023a; Sun *et al*., 2023b)

#### Drivers of infection prevalence analysis

Bio15 may explain the prevalence of ranavirus (AICc = 51.31 and AICc weight = 0.25). Bio15 stands for Precipitation seasonality, which is defined as the measure of the variation in monthly precipitation totals over the course of the year. Our model shows that the prevalence may become higher during the dry summer and become lower during the rainy seasons (bio15). It is followed by bio08, bio18, elevation, and life stage (Table S2 and Supplementary Fig S4).

#### Species distribution modeling using MaxEnt

The Maxent distribution model reported an AUC value of 0.85±0.01 providing fairly high robustness to the model. According to the model, the distribution of ranaviruses in GAR had higher responsiveness towards the variables bio2, bio3, bio5, bio11, bio16, bio17 and bio18. The model predicted that suitable habitats for ranaviruses prevail in southwestern, southeastern, central and northeastern regions of Guangxi (Fig7 a).

#### The influence of land use factors on ranavirus presence

Infections are recorded from both natural environments and modified environments by humans. It has been recorded inside forests, in the vicinity of villages, rivers/canals, reservoir/ponds and other construction areas. It appears that there is no barrier for the disease transmission (Fig 7 b)

**Figure 7.**
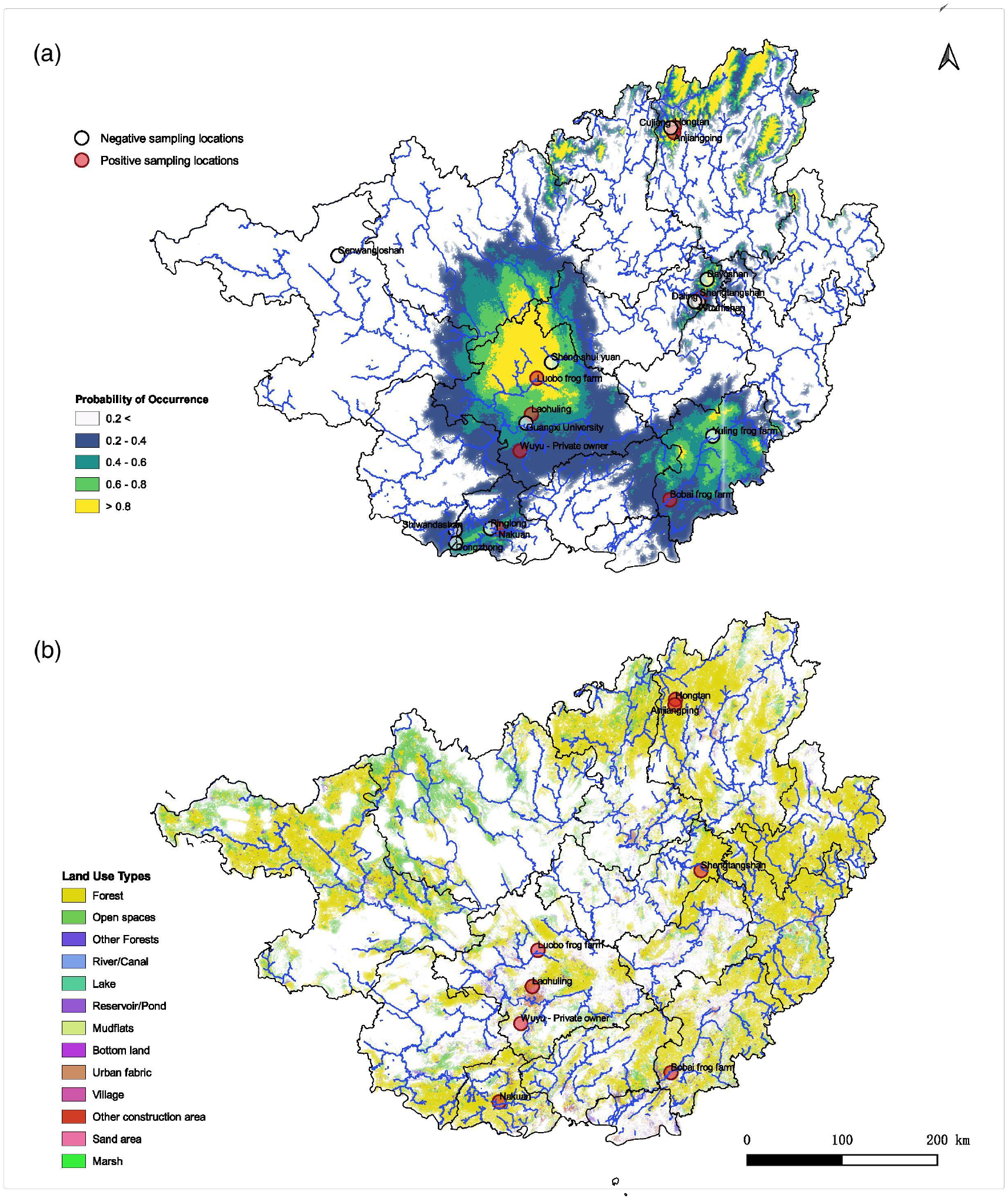
Predicted habitat suitability of ranaviruses in Guangxi region and its association with land use patterns. (a) demonstrates the probabilities of distributions in GAR with higher probability of occurrence towards the central, northeastern and southeastern regions of the map b) Spatial data map of the GAR representing how different land use patterns may contribute to the presence of ranavirus cases according to the geographic location. Forest habitats as well as urban settings seem to be associated with the predicted distribution of ranaviruses in GAR.

#### Native species infected and threat to biodiversity conservation

The emergence of infectious diseases with a broad host range has become one of the main threats to biodiversity as they can have a dramatic impact on entire communities. This is very concerning especially in small populations where recovery is slow as the ability of crossing species barriers of ranavirus can give rise to catastrophic consequences (Earl and Gray, 2014; Price *et al*., 2014). It has indicated up to 80% declines among ranavirus die-off sites among in common frog abundance in England (Teacher *et al*., 2010). In addition, amphibian recruitment attenuated in consecutive years signifying poor recovery following population declines at sites where ranavirus die-offs occurred (Petranka *et al*., 2003). There were 14 native species infected with ranavirus (11 species of anurans, 2 species of fish and 1 species of testudines) (Table S2). Out of 11 species of anurans, 9 species belonging to families of Ranidae, Rhacophoridae, Microhylidae and Hylidae are considered Least concern (LC) in IUCN Threat levels. Both species belonging to the family Dicroglossidae are known to be vulnerable (Table S2). Northern snakehead (*C. argus*) and large-scale loach (*P. dabryanus*) are widely used as cultured fish species and northern snakehead is considered to be Least concern (LC). However yellow pond turtle (*M.mutica*) which is widely used in pet industry considered as Critically endangered according to IUCN criteria. The individuals available at the pet markets are captive bred.

## DISCUSSION

### Presence of the ranaviruses in GAR

#### Epidemiology of ranavirus

Our study shows that ranaviruses are present across all classes of ectothermic vertebrates in south China, spanning various natural habitats and aquaculture facilities, highlighting the extensive reach of this disease across the region and possibly in other parts of Asia. The high number of infected amphibian species (14 in total), including both native and introduced host species, underscores the importance of monitoring and managing the spread of ranaviruses, particularly involving the ranaculture, aquaculture, mariculture and the pet trade. We also observed that susceptibility to ranaviruses varies greatly among host species, which is consistent with the findings from previous research (Bruner *et al*., 2015). The variation in infection prevalence among anuran families and the greater susceptibility of aquatic species compared to semi-aquatic, terrestrial, and arboreal species, suggest that host ecology is an important determinant of ranavirus disease dynamics. Our findings also underscore the need to consider multiple factors when investigating ranavirus outbreaks. Shedding rates, behavior, community composition, and interspecific variation in susceptibility are all likely to influence the likelihood, dynamics, and outcome of ranavirus outbreaks (Bruner *et al*., 2015).

The results of this study advance our understanding of the true burden of ranavirus infections in south China and highlight the potential threats it poses to its biodiversity, frog farming, and the pet industry. Prior to this research, only one case of ranavirus infecting cultured hybrid grouper had been recorded in GAR (Xiao *et al*., 2019). The high number of infected individuals and species found in this study suggests that GAR is a “burden hotspot,” in the context of Lessler *et al.,* (2017).

These findings are of significant concern given the high biodiversity of GAR and its location within the Indo-Burma biodiversity hotspot (Myers *et al*., 2000). Furthermore, the proximity of GAR to Yunnan province, an amphibian hotspot in terms of high species diversity that is already threatened (Chen and Bi, 2007), is also of concern. The transmission of ranaviruses in these regions could have severe consequences for regional biodiversity.

Ranavirus disease have been documented in various native wild species and cultured species in China (Herath et al., 2021), including the Critically Endangered (IUCN threat categories) Chinese giant salamanders (Chen *et al*., 2013). High mortality rates have been recorded in some cases, such as the 90% mortality in black-spotted pond frogs (*R. nigromaculata*) tadpoles (Mu *et al*., 2018; Yu *et al*., 2020). However, we did not witness mass mortality during the course of our study.

Given the pervasiveness of the disease, we emphasize the need for increased screening efforts to address the risks of ranavirus infections due to intensive aquaculture, ranaculture, and mariculture, as well as the pet industry.

#### Phylogenetic analysis based on the major capsid protein (MCP) gene

As the MCP gene is highly conserved, it is making it a desirable region to target for identifying the presence of ranavirus. Our MCP gene analysis identified only two strains of ranaviruses, Rana nigromaculata ranavirus and tiger frog virus, in the Guangxi region. Rana nigromaculata ranavirus strain that we found is identical to each other and to other ranaviruses reported from China, Japan, and Korea (Huang *et al*., 2009; Mu *et al*., 2018; Lei *et al*., 2012; Une *et al*., 2009; Yu *et al*., 2020; Zhang *et al*., 2001). The sequencing attempt on the ranaviruses detected in fish and reptiles was futile and not included in the phylogenetic analysis. The limited Ranavirus lineage diversity suggests a recent introduction, with rapid spread among different taxa due to their broad host range and transmission facilitated by regional and international trade of introduced species such as the American bullfrog and tiger frog (Both *et al*., 2011; Mazzoni *et al*., 2009; Sriwanayos *et al*., 2020).

Our results point towards interspecies and interclass transmission of these ranaviruses. Ranaviruses are known to infect fish, amphibians and reptiles, presumably due to host-switching events. The group of newly acquired genes in the ranavirus genome may have undergone recent adaptive changes that have facilitated interspecies and interclass host switching (Abrams *et al*., 2013). A phylogenetic analysis indicates that Rana nigromaculata ranavirus infects both introduced farmed tiger frogs and native frog species, with a considerable number of cases of the disease in farmed frogs. Previous studies also have shown the transmission of ranaviruses to native species from cultured species (Cunningham *et al*., 2015; Zhou *et al*., 2013; Chen *et al*., 2013). This suggests that the disease may have initially been introduced through cultured species.

So far, Asia harbors four out of seven ranavirus species recognized by the International Committee on Taxonomy of Viruses (Chinchar *et al*., 2017). Common midwife toad virus, frog virus 3, Santee-Cooper ranavirus and Singapore grouper iridovirus have been recorded in Asia while ambystoma tigrinum virus, epizootic haematopoietic necrosis virus and European North Atlantic ranavirus (or their isolates) have not yet been reported from the region (Herath *et al*., 2021).

Disease transmission may have been facilitated by poor biosecurity measures in GAR and throughout Asia (Herath *et al*., 2021; Sriwanayos *et al*., 2000). The possibility of interspecies transmission is supported by the close proximity of many main and sporadic sampling sites to frog farms, markets and pet markets which are well-established throughout GAR. The possibility of interclass transmission is supported by the presence of the disease in species such as tiger frogs, large scale loach, northern snakehead, yellow Pond Turtle, and golden albino axolotl, which are kept in adjacent containers for sales.

Our results point towards the potential role of fish and reptiles as reservoirs for ranavirus, given their ability to harbor subclinical infections (Brenes *et al*., 2014). These subclinical infections could contribute to the persistence of the pathogen in the environment, particularly when highly susceptible hosts, such as amphibians that rely on cool-wet conditions, are absent due to seasonal fluctuations in temperature and rainfall.

#### Co-infection with ranavirus and *Bd*

Another significance of our results is the identification of co-infection of one population with ranavirus and *Bd*. This is likely the first record of such co-infection in Asia. Previous studies have reported co-infection cases in other regions, including South America and Turkey (Warne *et al*., 2016; Erişmiş *et al*., 2019). The co-infection of ranaviruses and *Bd* is of concern, as it can lead to high mortality and morbidity. This happens through a primary infection with one pathogen weakening the immunity of the host, making it more susceptible to secondary infections (Herczeg *et al.,* 2021). This is particularly alarming in the context of amphibian populations already facing stress from factors such as climate change and anthropogenic stressors.

#### Climatic niche analysis

Our climatic niche analysis shows that the ranaviruses in our study are capable of adapting to a wide range of climatic conditions. The adaptability, along with the high number of asymptomatic cases and low mortality rates observed, suggests that these ranaviruses can persist and spread sub-clinically throughout the region without drawing much attention (Gray and Bruner, 2015).

The niche modeling further indicates that precipitation seasonality plays an important role in ranavirus prevalence in GAR. With climate change leading to rising temperatures, the spread of ranaviruses may be facilitated, potentially causing mass die-offs in the region. Our spatial analysis also sheds light on the transmission pathways and habitats of the ranaviruses. The detection of ranavirus in a variety of habitats, including forests, villages, rivers/canals, reservoirs/ponds, and construction areas, suggests a potential transmission pathway among ectothermic vertebrate classes, which has not been previously reported.

The presence of ranavirus in wildlife in nature reserves or forest areas is particularly concerning, as it demonstrates that the disease has spread even into protected areas. With no effective treatment currently available to reduce mortality and morbidity in wild populations (OIE, 2019), strict control measures, such as limiting international animal trade and implementing disease screening, must be followed.

## Conclusion

In conclusion, our study provides compelling evidence that ranavirus disease is widespread throughout GAR. With infection prevalence rates reaching as high as 100% in some wild frog species and even penetrating nature reserves, the current situation is of concern. Our research highlights the common presence of infections in culture facilities and pet farms, which likely serve as primary sources for the movement and transmission of ranaviruses throughout the region, country, and across the world through animal trade. Our findings suggest a recent introduction of ranaviruses to the GAR, followed by rapid transmission across various habitats. The co-infection of ranaviruses and *Bd* adds an extra layer of complexity to disease management, making it increasingly challenging to address. To mitigate the risk and impact of these pathogens, we strongly recommend implementing well-planned, systematic surveillance throughout Asia and enforcing stringent biosecurity measures to control further transmission.

## Acknowledgements

We thank Guangxi University Laboratory Startup Funding (MM), China Scholarship Council Fellowship for graduate studies. We are also grateful to staff members of nature reserves for granting permission and assisting with fieldwork. Further owners and staff members of the pet markets, frog farms and private collections are also acknowledged. We also express our gratitude Prof. Thomas B. Waltzek (Washington State University), Prof. Ellen Ariel (James Cook University) and Prof. Jess Brunner (Washington State University) Prof. Kun Fang, Cao (Guangxi University), Prof. Aiwu, Jiang (Guangxi University), Prof. Christos Mammides (Guangxi University) and all the lab members of Ecology, Development and Evolutionary biology (EED) of Guangxi University.

## EXTENDED MATERIAL

**Table S1:**
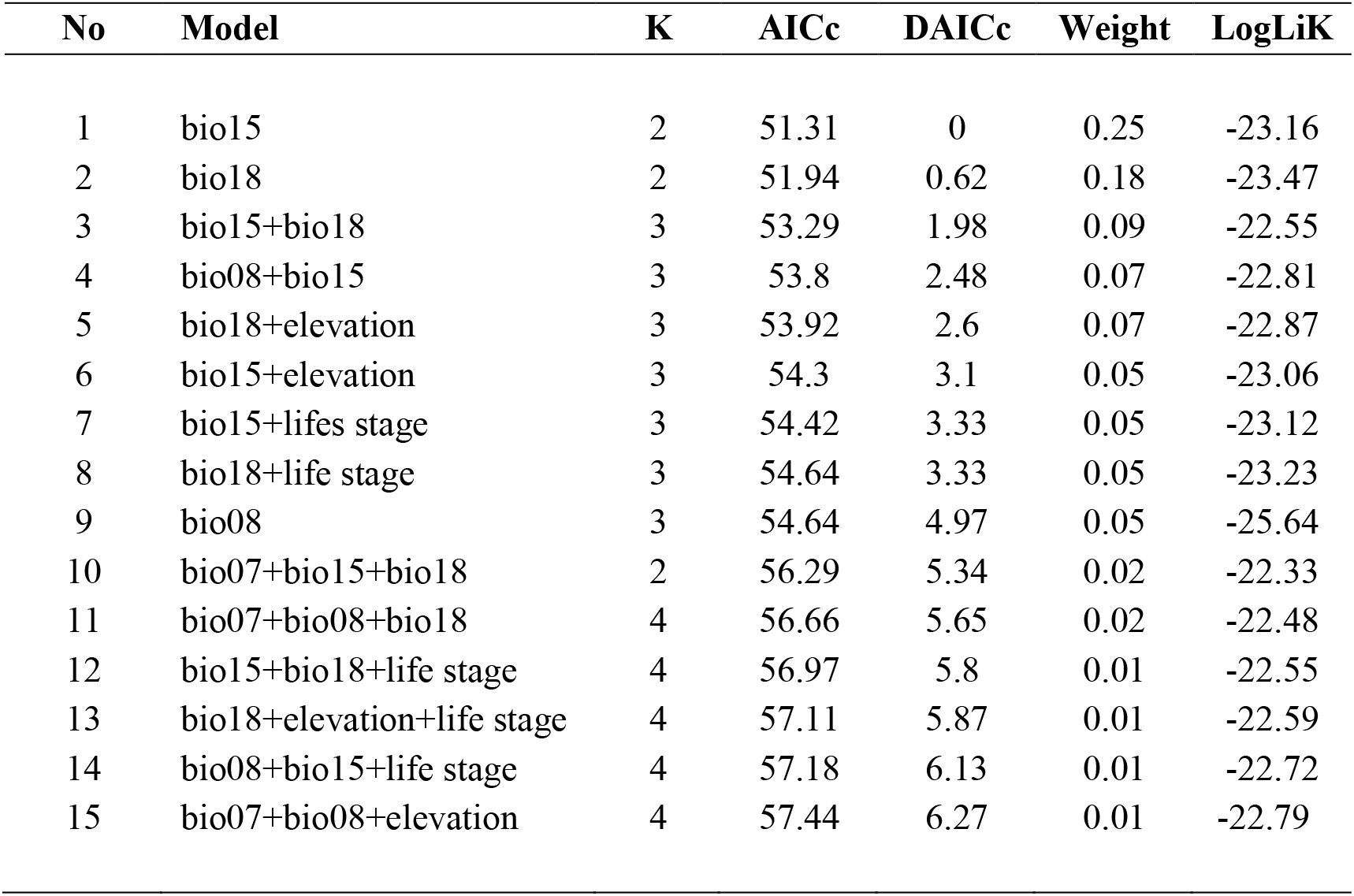
Model used for the analysis. This includes different possible combinations.

| No | Model | K | AICc | DAICc | Weight | LogLiK |
| --- | --- | --- | --- | --- | --- | --- |
| 1 | bio15 | 2 | 51.31 | 0 | 0.25 | -23.16 |
| 2 | bio18 | 2 | 51.94 | 0.62 | 0.18 | -23.47 |
| 3 | bio15+bio18 | 3 | 53.29 | 1.98 | 0.09 | -22.55 |
| 4 | bio08+bio15 | 3 | 53.8 | 2.48 | 0.07 | -22.81 |
| 5 | bio18+elevation | 3 | 53.92 | 2.6 | 0.07 | -22.87 |
| 6 | bio15+elevation | 3 | 54.3 | 3.1 | 0.05 | -23.06 |
| 7 | bio15+lifes stage | 3 | 54.42 | 3.33 | 0.05 | -23.12 |
| 8 | bio18+life stage | 3 | 54.64 | 3.33 | 0.05 | -23.23 |
| 9 | bio08 | 3 | 54.64 | 4.97 | 0.05 | -25.64 |
| 10 | bio07+bio15+bio18 | 2 | 56.29 | 5.34 | 0.02 | -22.33 |
| 11 | bio07+bio08+bio18 | 4 | 56.66 | 5.65 | 0.02 | -22.48 |
| 12 | bio15+bio18+life stage | 4 | 56.97 | 5.8 | 0.01 | -22.55 |
| 13 | bio18+elevation+life stage | 4 | 57.11 | 5.87 | 0.01 | -22.59 |
| 14 | bio08+bio15+life stage | 4 | 57.18 | 6.13 | 0.01 | -22.72 |
| 15 | bio07+bio08+elevation | 4 | 57.44 | 6.27 | 0.01 | -22.79 |

**Table S2:** Infected native species with their life stage and IUCN conservation status. Conservation status is based on IUCN categories with acronyms representing from low to high extinction risk: LC, least concern; NT, near threatened; VU, vulnerable; EN, endangered; DD, Data Deficiency

| Family | Species | Infected life stage | IUCN status |
| --- | --- | --- | --- |
| Ranidae | *Amolops ricketti* | Adults | LC |
|  | *Amolops chunganensis* | Adults | LC |
|  | *Odorrana gramine* | Adults | LC |
|  | *Odorrana versabilis* | Adults | LC |
|  | *Rana hanluica* | Adults | LC |
|  | *Hylarana guentheri* | Tadpoles | LC |
| Dicroglossidae | *Quasipaa spinosa* | Adults | VU |
|  | *Quasipaa boulengeri* | Adults | VU |
| Hylidae | *Hyla simplex* | Tadpoles | LC |
| Microhylidae | *Microhyla pulchra* | Tadpoles | LC |
| Rhacophoridae | *Polypedates megacephalus* | Tadpoles | LC |
| Cobitidae | *Paramisgurnus dabryanus* | Adults | - |
| Channidae | *Channa argus* | Adults | LC |
| Geoemydidae | *Mauremys mutica* | Adults | CR |

**Figure S1:**
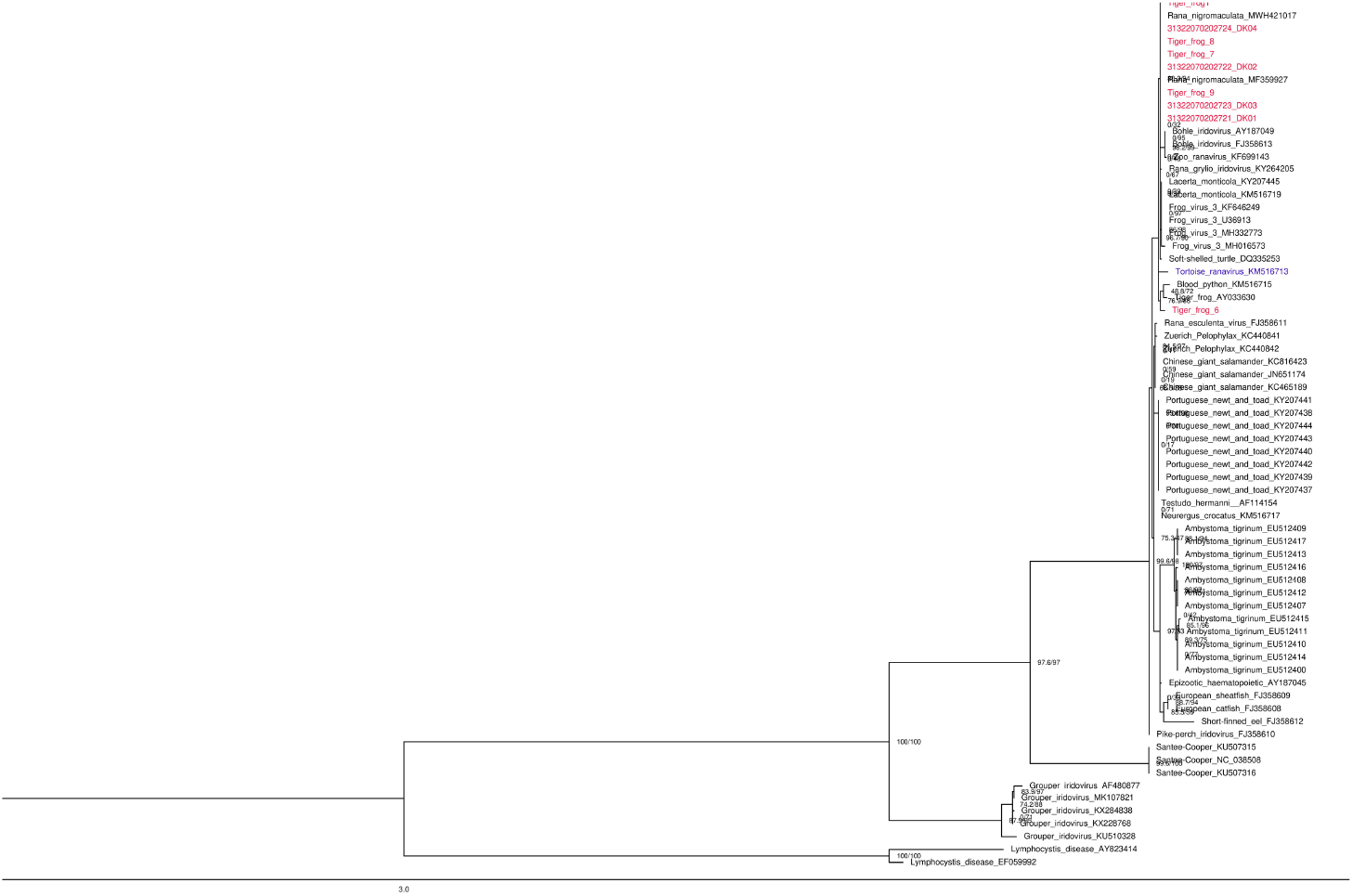
Phylogenetic tree based on the MCP gene sequences made using IQ-TREE.

**Figure S2:**
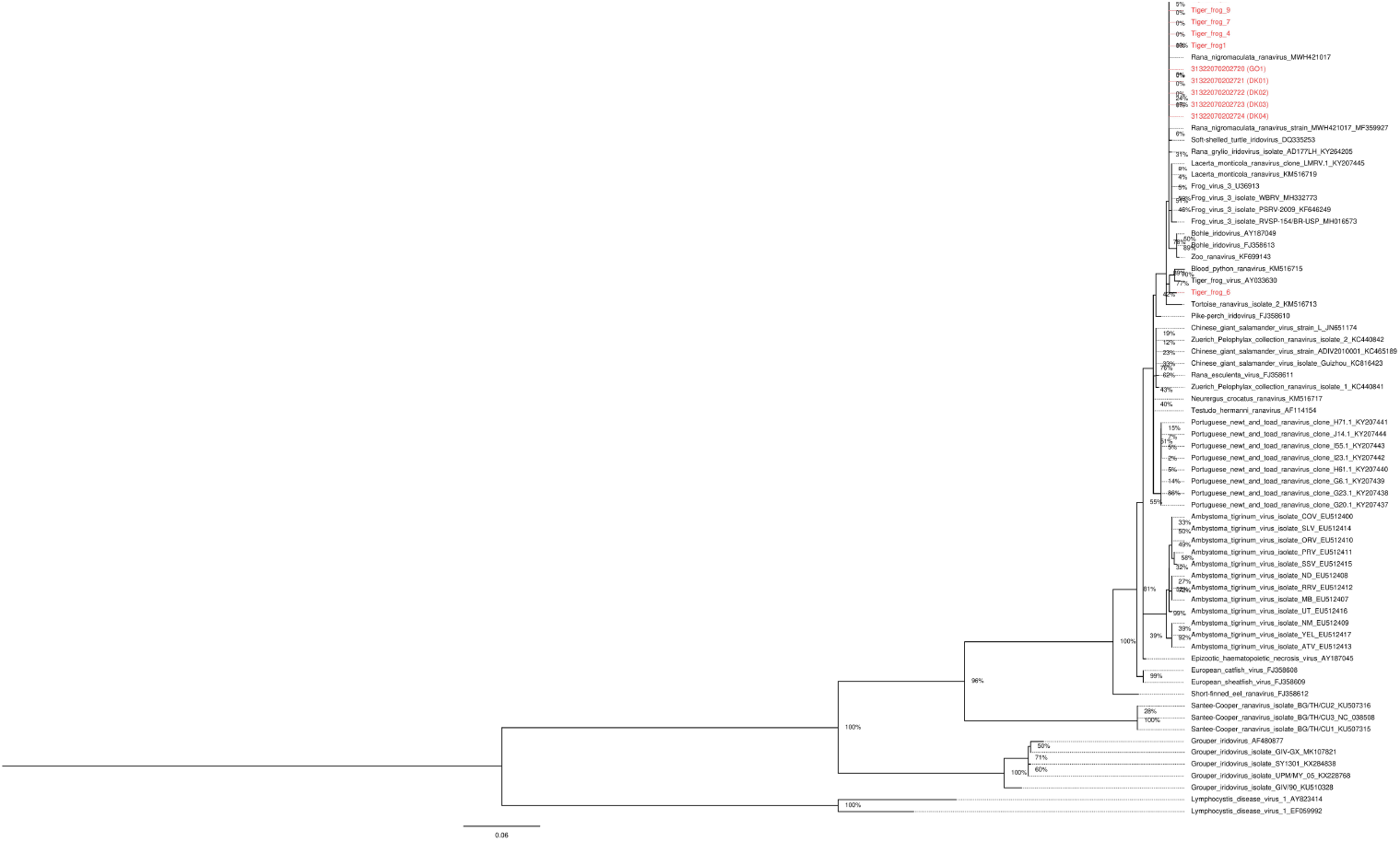
Phylogenetic tree based on the MCP gene sequencs made using MEGA.

**Figure S3:**
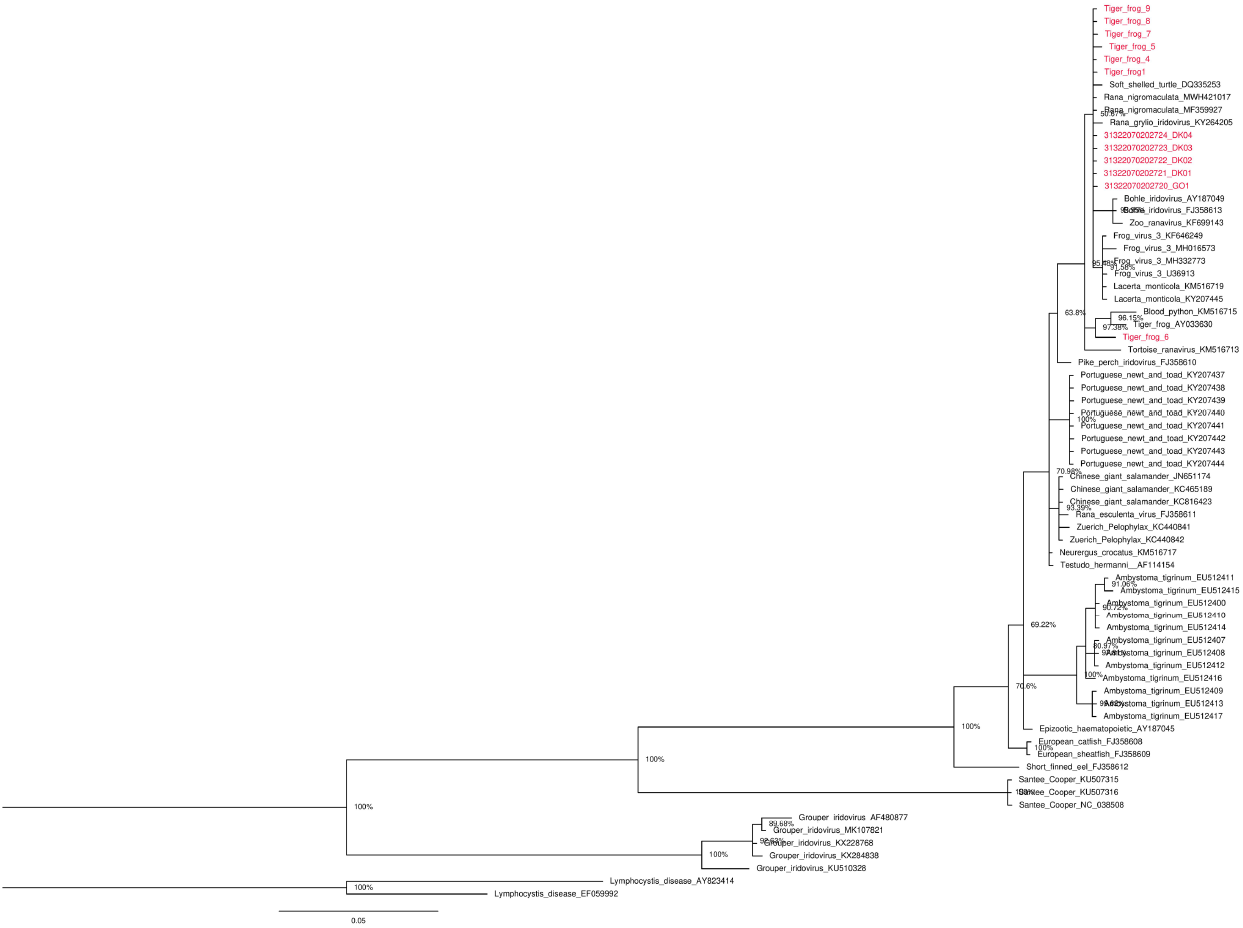
Phylogenetic tree based on the MCP gene sequences made using MrBayes.

**Figure S4:**
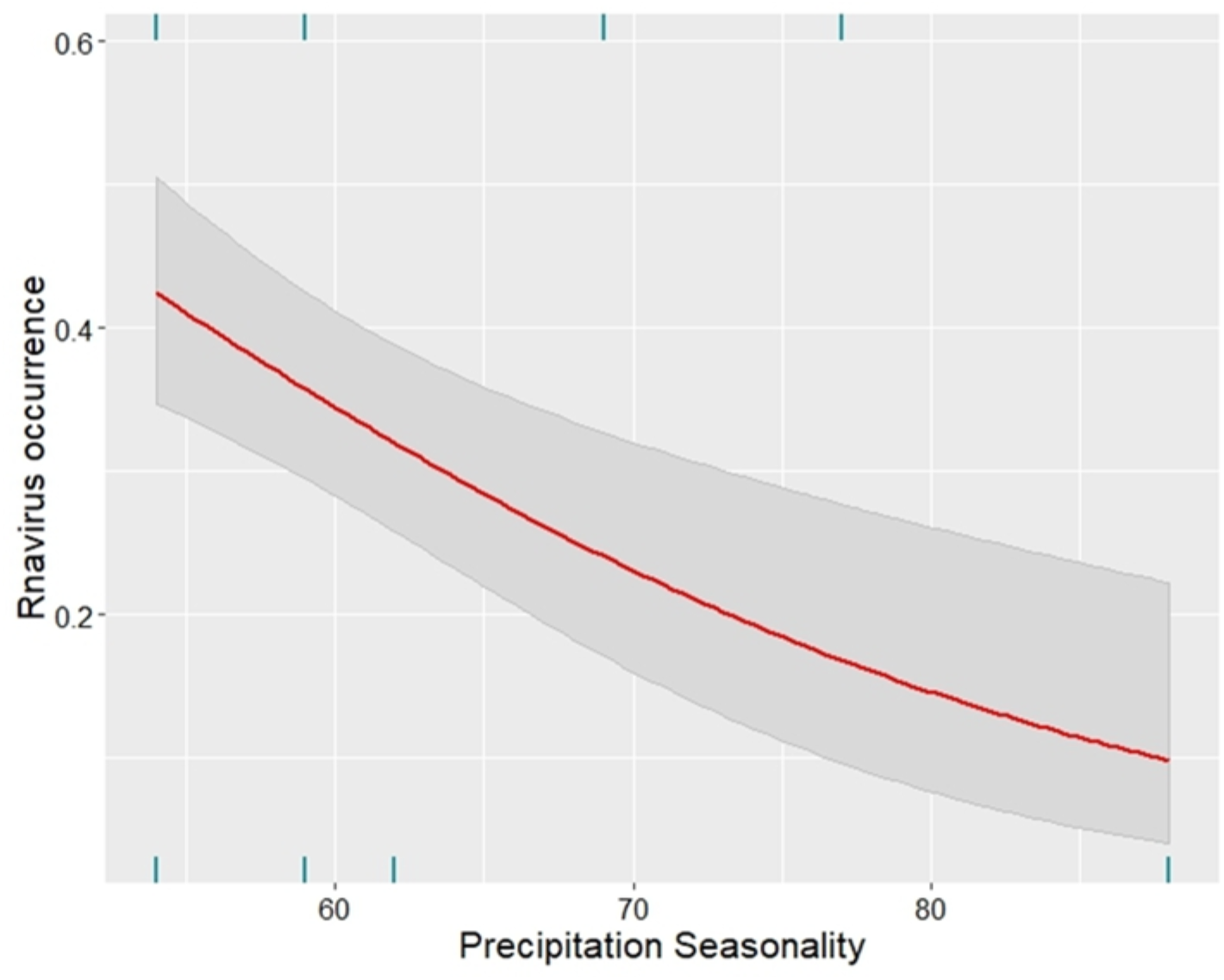
Precipitation seasonality.

**Figure S5:**
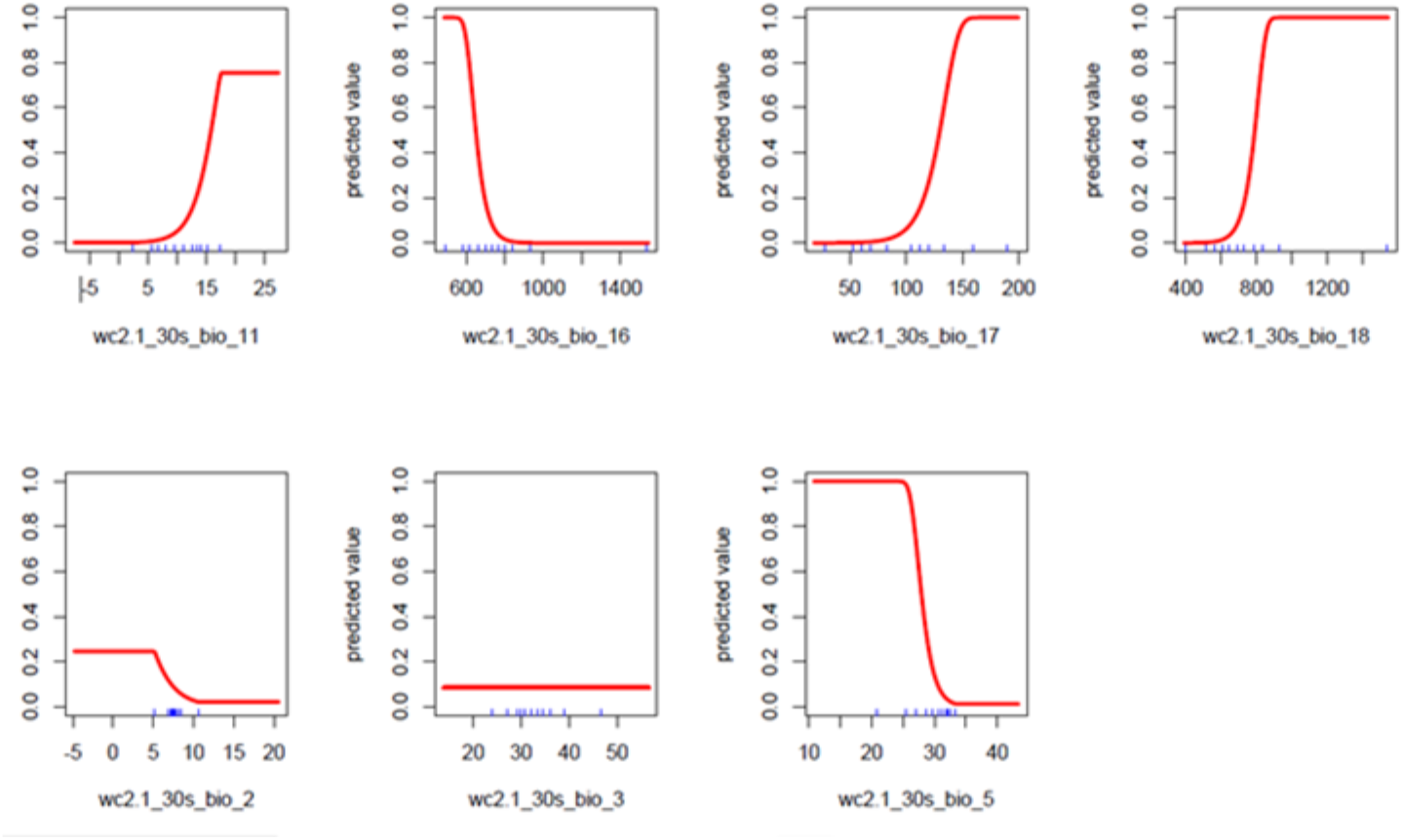
Response curves.

## Notes

### Competing Interest Statement

The authors have declared no competing interest.

